# Weak neural signatures of spatial selective auditory attention in hearing-impaired listeners

**DOI:** 10.1101/675595

**Authors:** Lia M. Bonacci, Lengshi Dai, Barbara G. Shinn-Cunningham

## Abstract

Spatial attention may be used to select target speech in one location while suppressing irrelevant speech in another. However, if perceptual resolution of spatial cues is weak, spatially focused attention may work poorly, leading to difficulty communicating in noisy settings. In electroencephalography (EEG), the distribution of alpha (8–14 Hz) power over parietal sensors reflects the spatial focus of attention (Banerjee et al., 2011; Foxe and Snyder, 2011). If spatial attention is degraded, however, alpha may not be modulated across parietal sensors. A previously published behavioral and EEG study found that, compared to normal-hearing (NH) listeners, hearing-impaired (HI) listeners often had higher interaural time difference (ITD) thresholds, worse performance when asked to report the content of an acoustic stream from a particular location, and weaker attentional modulation of neural responses evoked by sounds in a mixture (Dai et al., 2018). This study explored whether these same HI listeners also showed weaker alpha lateralization during the previously reported task. In NH listeners, hemispheric parietal alpha power was greater when the ipsilateral location was attended; this lateralization was stronger when competing melodies were separated by a larger spatial difference. In HI listeners, however, alpha was not lateralized across parietal sensors, consistent with a degraded ability to use spatial features to selectively attend.

## Introduction

Knowing where to attend is often helpful when trying to communicate in noisy environments (Kidd et al., 2005). However, if an individual has difficulty perceiving spatial differences among competing sound sources, then they may have difficulty deploying spatial attention. Hearing-impaired (HI) individuals often report difficulty holding conversations in noisy environments, even when using hearing aids (Marrone et al., 2008a). These difficulties are likely arise from poor encoding of sound features in the auditory periphery (Shinn-Cunningham, 2008; Shinn-Cunningham and Best, 2008). Specifically, if the perceptual representation of the spectro-temporal structure of sound is degraded (Buss et al., 2004; Strelcyk and Dau, 2009), then higher-order features that arise from these local features, like pitch and location, will also be degraded. Since such features support source segregation and source selection, a degraded peripheral representation can lead to failures on tasks requiring attention to be focused on a source in a complex scene (Shinn-Cunningham and Best, 2008).

Neurophysiological correlates of selective attention are often obtained using electroencephalography (EEG). In particular, growing evidence suggests that the distribution of alpha (8–14 Hz) oscillatory power across parietal sensors reflects the spatial focus of attention, with alpha power increasing ipsilateral to the location being attended (Foxe and Snyder, 2011; Banerjee et al., 2011). It is thought that this increase in alpha reflects suppression of the representation of distractors in the contralateral location (Worden et al., 2000; Foxe and Snyder, 2011; Banerjee et al., 2011). If spatial attention is degraded, however, then these neural correlates of attention may also be degraded. In a recent study, we found that HI individuals were less sensitive to interaural time differences (ITDs, a key feature when determining the perceived direction of sound) than normal-hearing (NH) individuals (Colburn, 1982; Dai et al., 2018). Since perceived spatial differences are crucial for deploying spatial attention, then HI individuals may depend less on spatial cues to segregate and select objects in a complex scene. If this is the case, then EEG correlates of spatial attention, including the distribution of alpha power, may not be strongly modulated with the locus of attention in HI listeners.

In healthy young listeners, attention strongly affects the magnitude of onset-evoked responses, including the N1, which arises between 100 and 150 ms after an onset event (e.g., the start of a note in a melody or a plosive sound in speech). In particular, N1 event-related potentials (ERPs) to events in a sound stream are larger when the stream is attended compared to when it is ignored. However, we previously found that this attentional modulation of the N1 is weaker in HI listeners than in NH listeners (Dai et al., 2018). Specifically, when NH subjects were directed to attend one of three simultaneous melodies, N1s to attended notes were larger than N1s to ignored notes; however, this difference was significantly reduced in HI listeners. Consistent with a previous study of NH listeners (Choi et al., 2014), we also found a direct correlation between the magnitude of the attentional modulation of N1 and the ability of individual HI and NH listeners to perform the spatial selective attention task. These results suggest that a degradation of attentional modulation reflects a degraded ability to selectively attend. These HI listeners also had higher audiometric, ITD, and audibility thresholds than NH listeners, supporting the idea that degraded feature representation contributes to failures of selective attention (Shinn-Cunningham, 2008; Shinn-Cunningham and Best, 2008).

Given these results, we wondered whether the degraded spatial selective attention abilities of HI listeners might also manifest in a weaker lateralization of alpha power over parietal EEG sensors. Specifically, based on previous work, we expected to find that the NH listeners in our study would display lateralized parietal alpha during auditory stimulus presentation. Furthermore, in that study, we tested conditions when the competing melodies were separated by either large or small differences in perceived lateral position. Based on other studies in our lab (Deng et al., 2017), we expected that for these NH listeners—who we thought would be able to deploy spatial attention effectively—the lateralization of alpha would be greater when the perceived spatial separation of competing melodies was large compared to when it was small. However, we hypothesized that HI listeners would show a reduction in or even a lack of alpha lateralization, reflecting a weaker deployment of spatial attention. To test these ideas, we reanalyzed the EEG data collected in our previously published study (Dai et al., 2018) of spatial selective attention in NH and HI listeners.

## Methods

Data were taken from the experiment previously published in (Dai et al., 2018). The subjects, stimuli, experimental paradigm, and data collection from this experiment are summarized below.

### Experimental Task and Stimuli

Subjects were presented with three simultaneous melodies: a distractor melody, a leading melody, and a lagging melody (Fig. 1A). For each trial, either the leading or lagging melody was the target and the distractor was always ignored. The three melodies were differentiated by the timing of their notes. The distractor melody started first, consisting of 4 notes with a duration of 919 ms and inter-stimulus interval (ISI) of 959 ms. A leading melody started 490 ms after distractor onset, and consisted of 5 notes with a duration of 624 ms and ISI of 664 ms. A lagging melody started 200 ms after the leading melody onset, and consisted of 4 notes with a duration of 728 ms and ISI of 768 ms.

**Figure 1.**
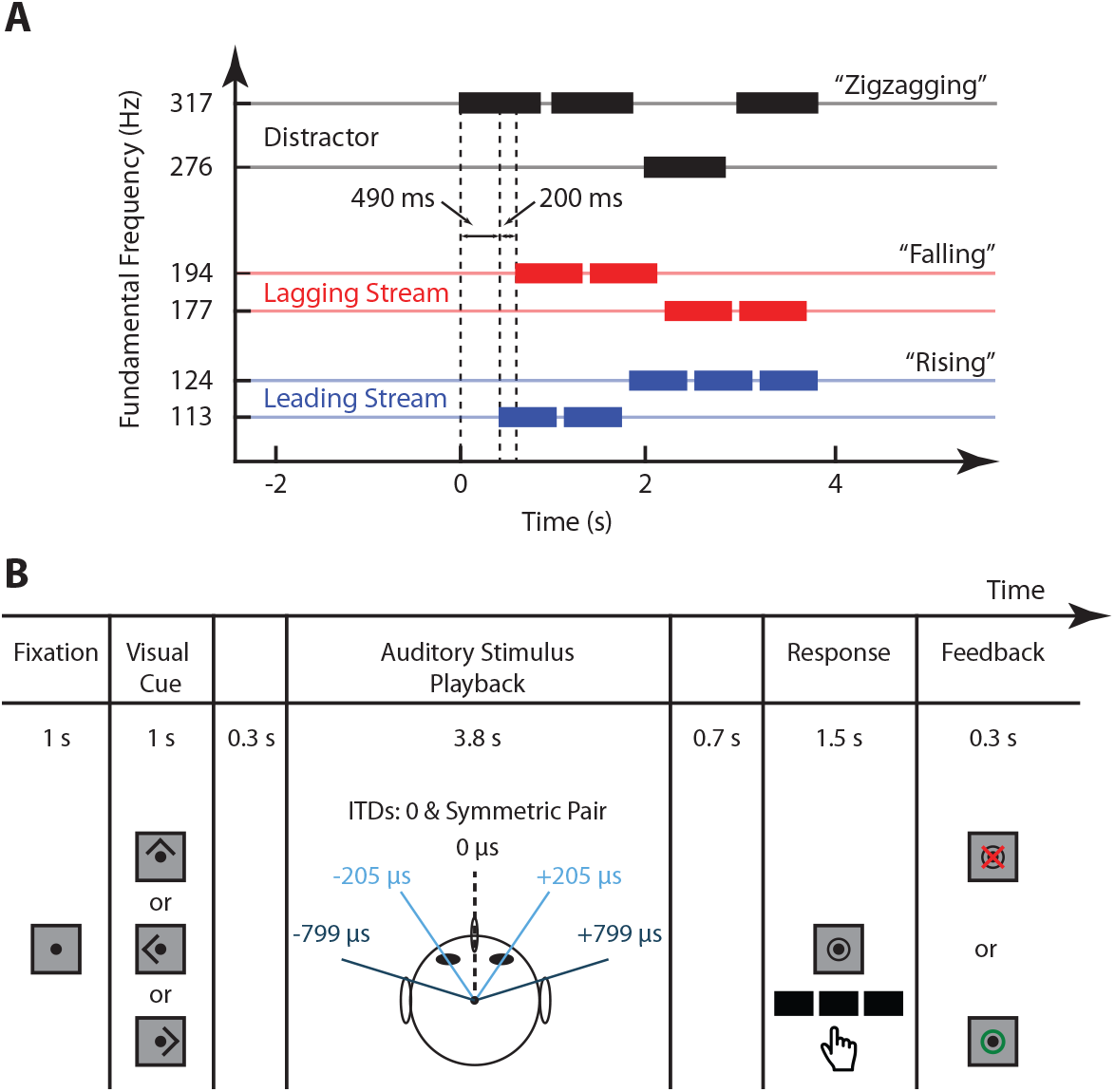
Stimuli and experimental paradigm (figure adapted from (Dai et al., 2018)). (A) Each trial presented three simultaneous melodies. Each melody consisted of a collection of high and low notes that formed one of three classes of pitch contours: “rising”, “falling”, or “zigzagging”. A distractor melody that was never the designated target always started first, with complex tones of fundamental frequency 276 Hz or 317 Hz. Next, the leading melody started, with each note having a fundamental frequency of either 177 Hz or 194 Hz. Finally, the lagging melody started, with fundamental frequencies that were either 113 Hz or 124 Hz. (B) For each trial, subjects fixated on a central point until a visual cue was given to either attend left, right, or center. On each trial, the correspondence between streams (distractor, leading, and lagging) and direction (left, right, and center) was selected randomly. Left and right melodies were spatialized using symmetric ITD pairs, and two conditions were tested: one with small ITDs (light blue) and one with large ITDs (dark blue). After stimulus presentation, subjects were asked to report the pitch contour of the cued melody and were given visual feedback on the correctness of their response.

Each of the three melodies was composed of high (H) and low (L) notes, differing in their fundamental frequency (F0). The H and L notes within a stream were relatively close in pitch, while the pitch separation *between* streams was larger. Distractor tones were comprised of a sinusoid of a fundamental (276 Hz or 317 Hz) and its first three harmonics, all at equal amplitude. Leading and lagging tones were broader band, consisting of a fundamental (leading F0: 113 Hz or 124 Hz, lagging F0: 177 Hz or 194 Hz) and its first 33 harmonics, all at equal amplitude. Notes were gated on and off with cosine-squared ramps of duration 10 ms and 100 ms for onset and offset, respectively.

Notes in each melody were arranged such that they formed a pitch contour that was “rising”, “falling”, or “zigzagging”. For “rising” melodies, the first note was low and transitioned to the high note for all subsequent notes (e.g., L-H-H-H). “Falling” melodies started high and transitioned to the low note for all subsequent notes (e.g., H-L-L-L). “Zigzagging” melodies started high or low, transitioned to the other note, and returned to the original note on the last onset (e.g., L-H-H-L or H-L-L-H). The pitch contour of each melody was chosen independently of the others, with each contour having equal probability (1/3).

The three melodies were spatialized such that one came from the left, one from the right, and one from center; the correspondence between melody and position was assigned randomly on each trial. Symmetric ITD pairs (either *±*205 *µ*s or *±*799 *µ*s) were used to spatialize left and right stimuli, while 0 *µ*s ITD was used for the center melody (Fig. 1B). Trials in which the symmetric ITD pair was small (*±*205 *µ*s) and those in which it was large (*±*799 *µ*s) were intermixed. All possible combinations of target location (left, right, or center), spatial separation (small or large ITD), and target stream type (leading or lagging) were tested, for a total of 480 trials (40 in each condition). Here, we only focus on attend-left and attend-right trials in each ITD condition, collapsed across all other conditions, as these should show the greatest alpha lateralization, providing the strongest test of alpha lateralization.

The structure of each trial is outlined in Fig. 1B. At the beginning of each trial, subjects fixated on a central dot for 1 s before a 1-s visual cue was given. The visual cue was an arrow that pointed left, right, or upward, signaling subjects to pay attention to the left, right, or center melody, respectively. After the visual cue, there was a 0.3-s quiet period, followed by 3.8 s of auditory stimulus playback, and another 0.7-s quiet period. Subjects were then prompted to identify the pitch contour of the cued sequence via button press. Subjects were given 1.5 s to respond after which visual feedback was given to indicate if the response was correct or not.

Training took place before testing to ensure that subjects could properly identify pitch contours of a single melody in quiet. This training consisted of two 12-trial blocks of a single stream. The first block tested leading streams (lowest F0), and the second block tested lagging streams (middle F0). Subjects were required to performed additional blocks until they achieved 8 of 12 correct trials for 7 consecutive blocks.

### Subjects

Data were collected from 25 NH listeners (13 male, 12 female, aged 20–52 years) and 15 HI listeners (8 male, 7 female, aged 20–59 years). All NH listeners had audiometric thresholds *≤* 20 dB hearing level (HL) at octave frequencies from 250 Hz to 8 kHz. HI listeners had bilateral symmetric sensorineural hearing loss. Audiometric thresholds for all HI listeners were *≥* 25 dB HL at one or more frequencies from 250–8,000 Hz, and threshold differences between the two ears were *≤* 20 dB at each frequency. NH and HI groups did not differ significantly in age (two-sided Wilcoxon Rank Sum test; rank sum = 329, P = 0. 5651) (Dai et al., 2018).

Stimuli were presented at 70 dB sound pressure level (SPL) for all NH listeners. For HI listeners, the level was adjusted, starting at 70 dB SPL and increasing in steps of 5 dB until a comfortable level was reached. Of the 15 HI listeners, 5 settled on 75 dB SPL while the remaining 10 settled on 70 dB SPL. These levels were used in training, prior to the testing. Therefore, given that all HI listeners were able to perform the melody contour identification task in quiet, audibility of the melodies was not a limiting factor in their performance.

All subjects gave informed consent before participating, and were compensated at an hourly rate and also paid a bonus of $ 0.02 for each correct response in order to maintain motivation. All procedures were approved by the Boston University Institutional Review Board.

### Data Collection

EEG data were recorded in 32 electrodes and sampled at 4096 Hz using the BioSemi ActiveTwo system along with its ActiveView acquisition software (BioSemi, Amsterdam, Netherlands). Scalp electrode positions were arranged according to the international 10-20 system, and two reference electrodes were placed on the mastoids. Event triggers were generated by MATLAB interfaced with Tucker-Davis Technologies System 3 (TDT, Alachua, FL) hardware and sent to the computer recording EEG data.

Subjects performed the experiment in front of an LCD monitor in a sound-treated booth. Stimuli were generated using MATLAB (MathWorks, Natick, MA) with the PsychToolbox 3 extension (Brainard, 1997). Sound stimuli were presented diotically via Etymotic ER-1 insert headphones (Etymotic, Elk Grove Village, IL) connected to Tucker-Davis Technologies System 3 (TDT, Alachua, FL) hardware which interfaced with the MATLAB software running the experiment. During the task, subjects were instructed to keep eyes open and positioned on a central fixation dot.

### Data Analysis

#### EEG Processing

Raw EEG data were first filtered from 1.5 to 50 Hz using a 6,000-point FIR band-pass filter. Data were then epoched and downsampled to 256 Hz before band-pass filtering again from 2–25 Hz. Scalp voltages were transformed to current source density (CSD) using CSD Toolbox (Kayser and Tenke, 2006). This transform has been shown to reduce spatial noise, which is useful when localizing alpha over parietal sensors (McFarland, 2015; Kayser and Tenke, 2015). No other artifact rejection was undertaken; all trials, whether correct and incorrect, were included in analysis. We chose this approach to ensure that we had a sufficient number of trials to analyze for all subjects.

#### Induced Alpha Power

To obtain the induced alpha response, we first removed phase-locked, or evoked activity. The evoked response was first calculated by averaging epochs across trials in each condition for each subject. This trial-average was then subtracted from each epoch to remove the phase-locked component for each trial. A short-time Fourier transform was then applied to each trial to estimate the power at each frequency in the alpha band (8–14 Hz). For each subject, an individual alpha frequency (IAF) was determined by finding the frequency in the range of 8–14 Hz whose magnitude was largest across cue-left and cue-right conditions in 10 parietal and occipital channels (P4, P8, PO4,O2, P3, P7, PO3, O1, Oz, Pz). Once an IAF was selected, power was extracted at this frequency to produce a single time series for each trial in each EEG channel.

For each subject, average alpha power over time was estimated for each condition using the median across all trials in that condition. First, attend leading and lagging trials were combined within each spatial attention condition (i.e., attend-left and attend-right). The median was then taken across the combined trials to estimate the average alpha power time series in each channel. The median was used instead of the mean in order to obtain an estimate that was robust to outliers, since no artifact rejection was performed. These trial-averaged time series were then normalized for each subject by dividing each time point by the average alpha power across time, sensors, and experimental conditions. Grand averages were obtained from these normalized time series. Quantities shown on topoplots represent averages across the stimulus period (0–3.14 s).

An attentional modulation index of alpha power, AMI_*α*_, was quantified for each subject, and is given by Eq. 1.

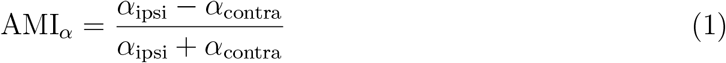

Note that *α*_ipsi_ is the average alpha power during the stimulus period, measured ipsilateral to the cued sequence (e.g., average alpha in left parietal channels during attend-left trials), and *α*_contra_ is this average alpha power, measured contralateral to the cued sequence, (e.g., average alpha in right parietal channels during attend-left trials). Positive values of AMI_*α*_ indicate that alpha power was overall larger when subjects attended the ipsilateral stimuli (i.e., the alpha response over a particular set of cortices was greater when ignoring the contralateral stimuli), as expected. Averages were calculated across left and right parietal and occipital channels separately, depending on the attention condition (i.e., left channels P3, P7, PO3, O1 for *α*_ipsi_ in attend-left trials and right channels P4, P8, PO4, O2 for *α*_ipsi_ in attend-right trials).

#### Significance Testing

We asked if there were differences in alpha modulation (AMI_*α*_) between NH and HI listeners in any of the ITD conditions tested. To determine if there were significant differences, we used a two-way mixed factors ANOVA, with the between-groups factor being hearing status (two levels: NH and HI) and the within-groups factor being ITD condition (two levels: small and large ITD). Kolmogorov-Smirnov tests of normality were conducted before obtaining ANOVA results. Tukey’s HSD was used *post hoc* to compare AMI_*α*_ between ITD conditions within each group (NH and HI). One-sample *t*-tests were also used *post hoc* to determine if AMI_*α*_ was significantly greater than zero. A two-way mixed factors ANOVA was also used to determine if differences in performance measures existed in the subset of trials reported here, which did not include attend center trials analyzed in (Dai et al., 2018). A *t*-test was used to determine if ITD thresholds were significantly different between NH and HI listeners, and Pearson’s method was used quantify the correlation between these ITD thresholds and performance measures.

## Results

### Behavior

#### HI listeners performed worse on the task, and had higher ITD thresholds than NH listeners

As we previously reported (Dai et al., 2018), HI listeners performed significantly worse on the spatial attention task than NH listeners. This result is summarized here by comparing the average percent correct scores, collapsed across attend-left and attend-right trials in each ITD condition (Fig. 2A). A two-way mixed ANOVA confirmed significant main effects of hearing status (*F* (1, 38) = 11057, *p* = 0.0001) and ITD condition (*F* (1, 38) = 186.05, *p* = 0.0003) on percent correct scores, and no significant interaction. Thus, HI listeners performed significantly worse on the task than NH listeners, and increasing the perceived spatial separation significantly improved performance overall.

**Figure 2.**
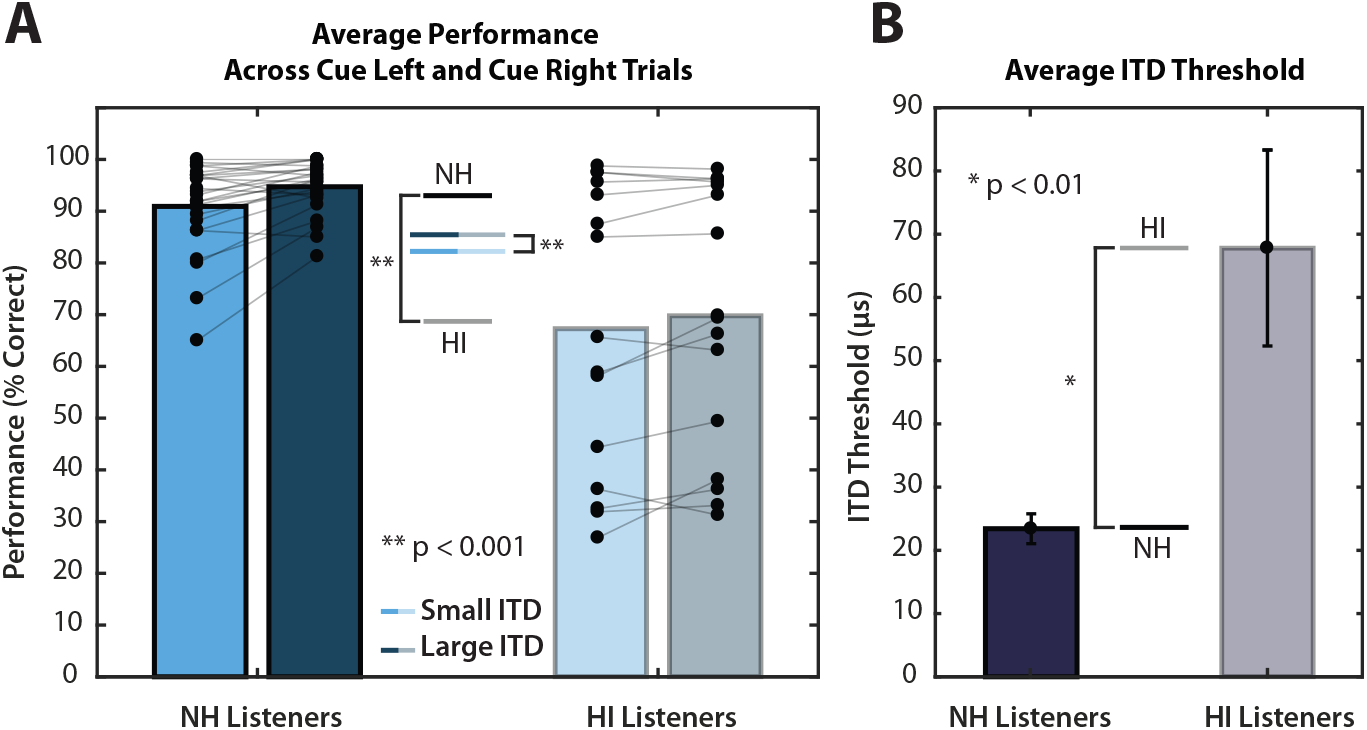
(A) Performance averaged across attend-left and attend-right trials. Asterisks indicate significant main effects of hearing status and ITD condition. (B) ITD thresholds for NH and HI listeners. HI listeners had significantly higher ITD thresholds than NH listeners (*p* < 0.01, *t*-test). Error bars represent the standard error of the mean.

Figure 2B shows ITD thresholds for NH and HI subject groups. ITD thresholds were measured separately for the leading, low pitch stimuli, and the lagging, high pitch stimuli. Since these thresholds were not significantly different within subjects for the two different stimuli, we averaged the two measured ITD thresholds for each subject. The average ITD threshold for NH listeners (23.46 *±* 11.79 *µ*s, mean *±* std. dev) was significantly lower than that for HI listeners (67.81 *±* 60.02 *µ*s, mean *±* std. dev) (*p <* 0.01, *t*-test). These results confirm that these HI listeners had significantly poorer spatial acuity than the NH listeners in our experiment. Previous results published in (Dai et al., 2018) found a significant correlation between average ITD threshold and average performance on the task for both NH and HI listeners. This correlation remained for the subset of data analyzed here, which excluded “attend center” trials; only trials for which the target was to the left or right were included (NH: *r* = *−*0.48*, p* = 0.0148, HI: *r* = *−*0.59*, p* = 0.029).

### Induced Alpha Power

#### In NH listeners, alpha was lateralized over parietal sensors, and this lateralization was stronger in the large ITD condition

Figure 3A shows the time course of induced alpha power averaged in left and right parietal-occipital sensors for NH listeners. In left sensors, alpha power was greater throughout the stimulus period (0–3.14 s) when subjects were cued to attend the melody on the left (blue trace), compared to when they were cued to attend the melody on the right (red trace). In right sensors, alpha power was greater throughout most of the stimulus period during attend-right trials. In comparing small and large ITD conditions, the difference in alpha power between red and blue traces appears to be larger in both left and right parietal-occipital sensors. In both conditions, we see that alpha power was modulated over time in NH listeners. After receiving a visual cue for where to attend, alpha decreased briefly and then increased before stimulus playback. Before the response period, alpha decreased again.

**Figure 3.**
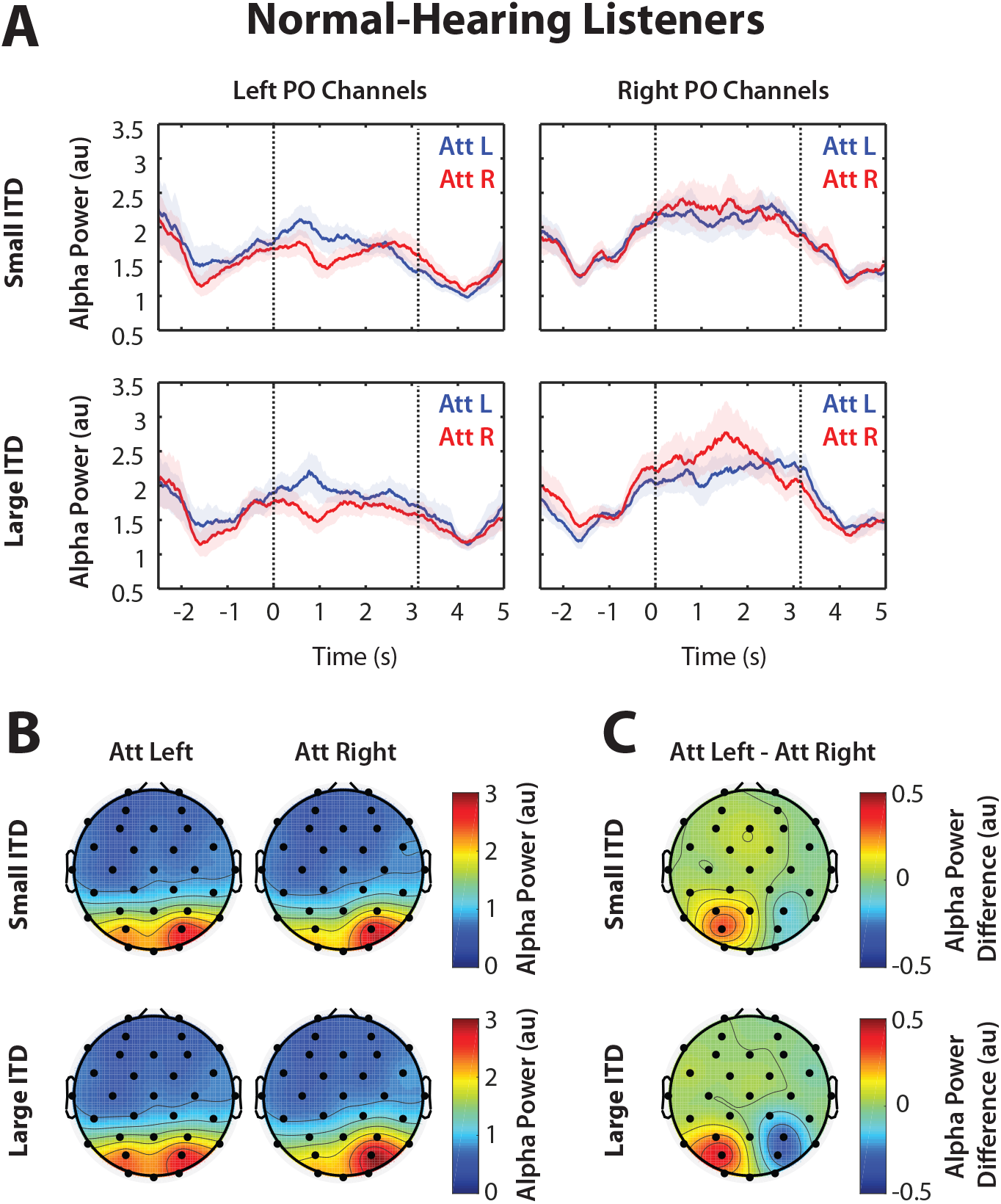
Grand average alpha power for NH listeners. (A) Grand average alpha power in left and right PO channels for both ITD conditions. Error bars represent the standard error of the mean. Dashed vertical lines specify the onsets of the first and last notes of auditory playback. (B) Grand average alpha power in 32 channels, averaged during the stimulus period (0–3.14 s). Average alpha is displayed separately for attend-left and attend-right trials. (C) Grand average alpha power differences between attend-left and attend-right trials during the stimulus period in each of the 32 channels.

Figure 3B shows alpha power averaged over the stimulus period (0–3.14 s) in each sensor for NH listeners. Here, we see an overall asymmetry, independent of the direction of attention: alpha power was always larger in right parietal sensors. This general asymmetry is consistent with the parietal spatial representation being asymmetric, and has been observed in other studies of alpha lateralization with spatial attention (Heilman and Van Den Abell, 1980; Pouget and Driver, 2000; Szczepanski et al., 2010; Ikkai et al., 2016). Importantly, however, in comparing attend-left and attend-right conditions, alpha was greater in left parietal sensors for attend-left trials compared to attend-right trials; in right parietal sensors, alpha was greater in attend-right trials. These differences are more apparent in Fig. 3C, where alpha is shown as the average difference between attend-left and attend-right trials during the stimulus period at each sensor on the scalp. Comparing small and large ITD conditions, we see that this alpha modulation with the direction of spatial attention was stronger when the perceived spatial separation was large.

#### In HI listeners, no alpha lateralization was observed across parietal sensors in either small or large ITD condition

Figure 4A shows the time course of alpha power averaged in parietal-occipital sensors for HI listeners. Unlike in NH listeners, alpha power in HI listeners did not appear to be modulated over time in either left or right sensors in either attention condition; alpha power did not even decrease after presentation of the visual cue as it did in NH listeners. There also appears to be no difference, in either set of parietal sensors, between attend-left and attend-right trials during the stimulus period. These results are similar between small and large ITD conditions.

**Figure 4.**
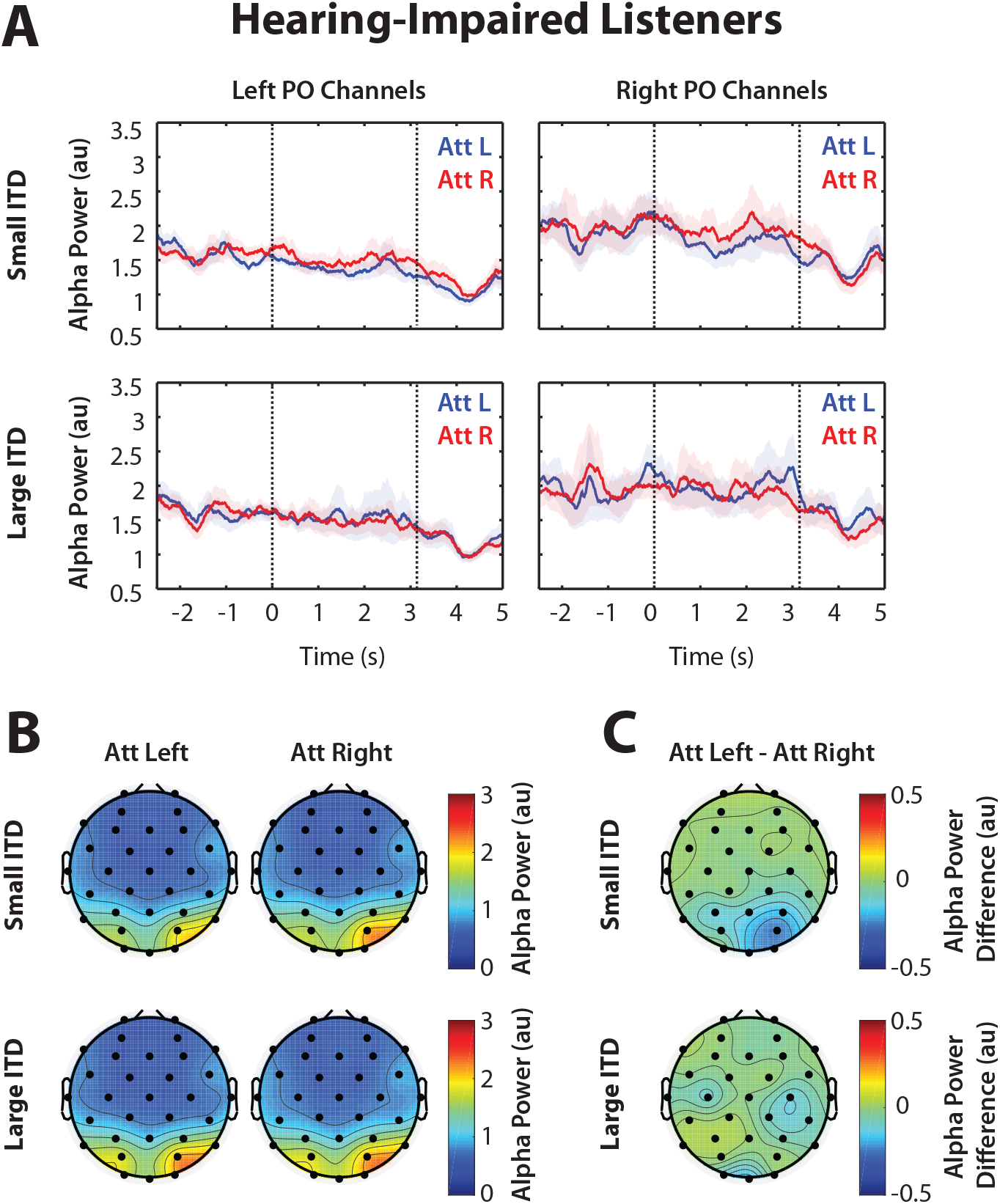
Grand average alpha power for HI listeners. (A) Grand average alpha power in left and right PO channels for both ITD conditions. Two subjects’ data were excluded from these time traces due to motion artifacts during the cue period. These subjects were only excluded in A since these artifacts occurred outside the time period of interest for subsequent analyses (0–3.14 s). Error bars represent the standard error of the mean. Dashed vertical lines specify the onsets of the first and last notes of auditory playback. (B) Grand average alpha power in 32 channels, averaged during the stimulus period (0–3.14 s). Average alpha is displayed separately for attend-left and attend-right trials. (C) Grand average alpha power differences between attend-left and attend-right trials during the stimulus period in each of the 32 channels.

Average alpha power during the stimulus period is shown in Fig. 4B. Here, we see the same asymmetry observed in NH listeners: greater alpha power in right parietal-occipital sensors than left sensors. However, unlike in NH listeners, there appears to be no substantial difference in any of these sensors between attend-left and attend-right trials (Fig. 4C). Assuming alpha lateralization is an indication that spatial features are being used for selective attention, these results suggest that HI listeners do not use these spatial features. Increasing the perceived spatial separation did not increase the amount of alpha modulation observed across parietal sensors.

#### There was a significant interaction between hearing status and perceived spatial separation

In order to characterize overall alpha modulation, we collapsed the alpha differences shown in Figs. 3C and 4C across parietal sensors that were mirrored across hemispheres, as shown in Fig. 5A. Here, alpha power is represented as the generalized difference between ipsilateral and contralateral attention conditions. In NH listeners, alpha power was greater in a particular set of parietal-occipital sensors when subjects were attending the ipsilateral sequence (i.e., ignoring the contralateral sequence). We found no difference between ipsilateral and contralateral attention conditions in HI listeners, however.

**Figure 5.**
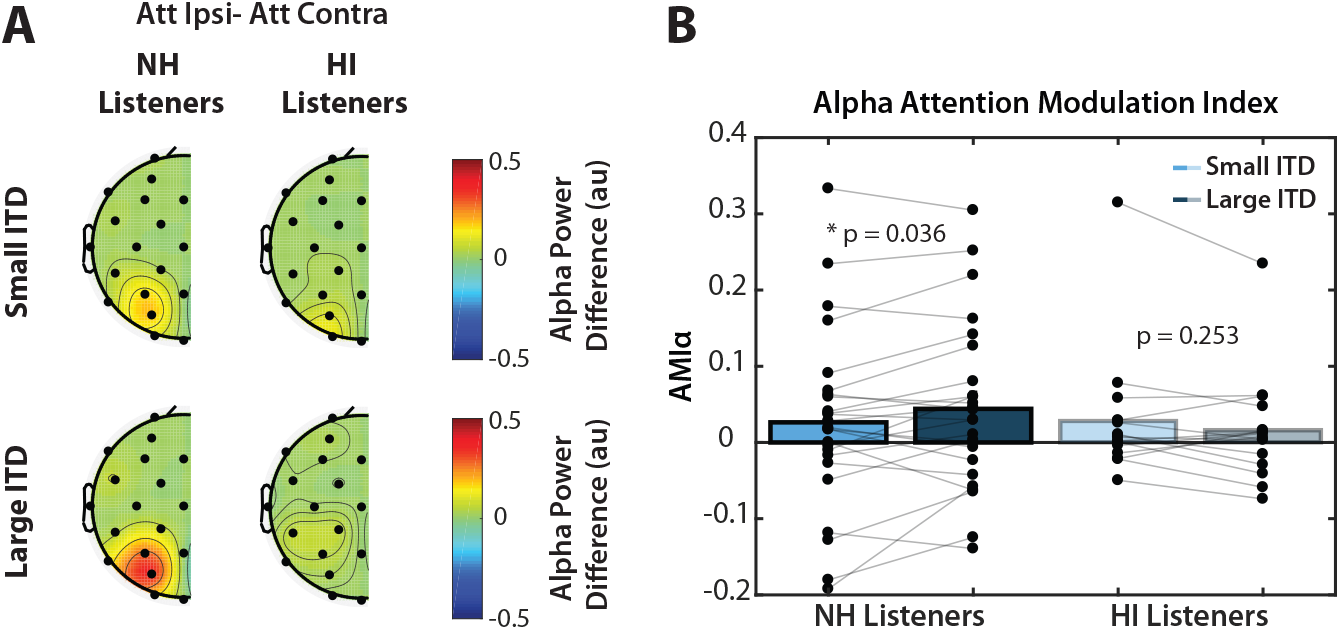
(A) Grand average alpha power differences between ipsilateral and contralateral attention conditions, mirrored across hemispheres. (B) Alpha Attention Modulation Index for each subject in small (light blue) and large (dark blue) ITD conditions. Asterisks indicate significant differences between ITD conditions at the *α* = 0.05 significance level (Tukey’s HSD). AMI_*α*_ was only significantly greater than zero for NH listeners in the large ITD condition (*p* = 0.028, *t*-test).

AMI_*α*_ was quantified for each subject and is shown in Fig. 5B. We asked whether there were significant differences in AMI_*α*_ between NH and HI listeners performing an auditory spatial attention task. The results of a two-way mixed ANOVA found no significant main effect of either hearing status (*F* (1, 38) = 0.157, *p* = 0.694) or ITD condition (*F* (1, 38) = 0.172, *p* = 0.681). However, there was a significant interaction between the two factors (*F* (1, 38) = 5.06, *p* = 0.0303). Tukey *post hoc* testing revealed that there was a significant difference in AMI_*α*_ between large and small ITD conditions in NH listeners (*p* = 0.036), but not in HI listeners (*p* = 0.253), suggesting that a larger perceived spatial difference contributes to greater alpha lateralization in NH listeners, but not in those with degraded spatial acuity. While it may initially seem surprising that there was no main effect of hearing status on alpha modulation, further analysis revealed that for NH listeners, AMI_*α*_ was significantly greater than zero for the large ITD condition (*p* = 0.028, *t*-test), but not for the small ITD condition (*p* = 0.137, *t*-test), whereas for the HI listeners, it was not significantly greater than zero in either condition. Thus, there is a floor effect on the results: alpha lateralization was only significant for the “best” listeners (the NH listeners) in the large ITD condition. This suggests that unlike NH listeners, HI listeners may not depend on spatial cues to maintain attention on the target stream even in the large ITD condition.

## Discussion

### To perform this task, listeners had to rely on spatial cues, at least initially

In our task, the target on a given trial could be either the leading or lagging stream, and the target could come from left, right, or center with equal likelihood. The leading, lagging, and distractor streams differed from one another consistently in their pitch and timing cues. However, the target was only defined by the visual cue for which direction to attend. Thus, all listeners had to initially use spatial information in order to select the target stream from the sound mixture.

All of our NH listeners performed well above chance in all conditions (Dai et al., 2018). While there were a few HI subjects who performed near chance in some spatial configurations (see Figure 2B in (Dai et al., 2018) for details), most performed well above chance levels. Thus, our results suggest that even in our HI group, most listeners were effective at using spatial attention to focus on the target melody, at least to some degree.

Once a target melody was the focus of attention, listeners could maintain attention on that target without using spatial information: the target always differed from the competing melodies in its pitch range and its note timing. In the current study, we do not have sufficient statistical power to explore the time course of alpha lateralization dynamics over the course of a trial. Instead, the current *post hoc* analyses considered sustained lateralization of alpha power over parietal EEG sensors, to test the hypothesis that sustained alpha lateralization would be weaker in HI listeners compared to NH listeners. However, if listeners transiently engage spatial attention and then maintain focus on a target using other features, it would not be reflected in our alpha lateralization metrics. Future experiments specifically designed to reveal such dynamics could lend more insight into how spatial attention is used by different listeners, and how this relates to their specific hearing acuity.

### Spatial acuity only predicts performance when spatial separations are near perceptual limits

A number of studies have demonstrated that HI listeners benefit less from spatial release from masking in multi-talker settings than do NH listeners (Marrone et al., 2008b; Srinivasan et al., 2016; Best et al., 2012), consistent with the current results. However, past studies linking spatial acuity measures with selective attention measures have produced conflicting results (e.g., (Strelcyk and Dau, 2009; Lőcsei et al., 2016)). In reconciling these discrepant findings, it is important to consider exactly what tasks are being used in a given study, and what is limiting performance.

If the spatial separation between competing sounds is large, even HI listeners with poor spatial acuity may be able to use spatial information effectively. For instance, one study of NH and HI listeners examined speech-in-noise performance when presenting two speech streams played with ITDs of *−*700 *µ*s and +700 *µ*s (an ITD difference of 1.4 ms) (Lőcsei et al., 2016). In this case, the large separation benefited both NH and HI listeners by roughly the same amount; moreover, ITD thresholds did not correlate with performance.

Another study (by the same research group in (Lőcsei et al., 2016)) found a significant correlation between ITD sensitivity and the ability to understand speech that is spatially separated from a target (Strelcyk and Dau, 2009). Importantly, in this study, the target was diotic and a single masker was played from left or right with an ITD of 740 *µ*s (about half the spatial separation between sources used in (Lőcsei et al., 2016)). When the spatial separation of sources is closer to the perceptual limit, it makes sense that ITD sensitivity is closely related to performance.

In our study, there were three competing streams that were separated by 799 *µ*s in the large ITD case (and by only 205 *µ*s in the small ITD case). Given that even our “large” ITD was smaller than that used in many studies, and given that our listeners heard a relatively complex scene with three concurrent streams, it is therefore not surprising that spatial acuity was directly related to the ability to perform the task.

Given these results, we did look to see whether the degree of alpha lateralization in an individual subject was related to their task performance. However, we found no such relationship. Importantly, our alpha lateralization metric only quantifies sustained spatial attention, so this is not particularly surprising. We suspect that in the right experiment, the strength of sustained alpha lateralization might be directly related to spatial attention performance; however, to see such effects likely requires an experiment in which the competing stream identities are confusable *except* in their spatial attributes.

### In NH listeners, parietal alpha lateralization likely reflects some combination of what location is the focus of spatial attention and how strongly listeners are sustaining spatial attention

Previous work has suggested that parietal EEG alpha lateralization reflects the locus of spatial attention; specifically, alpha power increases over sensors contralateral to ignored stimuli (Worden et al., 2000; Kerlin et al., 2010; Foxe and Snyder, 2011; Banerjee et al., 2011; Händel et al., 2011; Payne et al., 2013; van Diepen et al., 2016; Wöstmann et al., 2016). While most of these studies have identified this lateralization during visual spatial attention (Worden et al., 2000; Foxe and Snyder, 2011; Händel et al., 2011; Payne et al., 2013; van Diepen et al., 2016), considerably fewer have addressed alpha as a correlate of auditory spatial attention (but see (Kerlin et al., 2010; Banerjee et al., 2011; Wöstmann et al., 2016)). Here, we provide additional evidence for parietal alpha lateralization as a correlate of auditory spatial attention. In NH listeners, we observed that mean alpha power was greater in a particular set of parietal sensors when subjects attended the ipsilateral melody (i.e., ignoring the contralateral melody) when ITDs were large.

NH listeners showed less alpha lateralization for small ITDs than for large ITDs, and the lateralization was only significant for large ITDs. This pattern may be explained by two (not mutually exclusive) effects.

First, in another study from our lab, we have observed that alpha lateralization increases the farther off midline a listener directs auditory spatial attention (Deng et al., 2017). If the magnitude of alpha lateralization scales with eccentricity of attention, it would produce greater alpha lateralization in large ITD trials than in small ITD trials. The lack of a significant effect in the small ITD condition thus might be simply a matter of statistical power: there may be a small lateralization of alpha that we do not have the power to observe in this study.

Second, we have observed that when other sound features, such as pitch differences, differentiate one sound stream from another more effectively than do spatial features, listeners rely on those non-spatial features to maintain attentional focus (Bonacci and Shinn-Cunningham, 2019). In the current study, small ITDs may have been less reliable than the pitch separations and timing regularities of the streams in maintaining attention, so that alpha lateralization averaged over the three seconds of stimulus presentation was not significant. In contrast, when the ITD separation was large, it may have been more reliable than the pitch cue for our NH listeners, leading to significant sustained alpha lateralization in these trials. Indeed, previous studies have shown that when there are redundant features, their relative strengths determine how much influence each has on attention to an ongoing sound stream: as one feature becomes relatively stronger in differentiating competing streams, that feature is more influential, and vice versa (Maddox and Shinn-Cunningham, 2012).

We therefore believe that the difference in NH listenersâĂŹ alpha lateralization for small and large ITD conditions comes about through some combination of two factors. Specifically, alpha lateralization seems to scale with the spatial eccentricity of the target, and listeners may rely more heavily on spatial cues for larger spatial separations than for smaller separations.

### In HI listeners, alpha was never significantly lateralized, suggesting that HI listeners do not rely strongly on spatial cues to maintain attentional focus

Our HI listeners showed no significant alpha lateralization, even in large ITD trials. As already reported (Dai et al., 2018), our HI listeners also had worse spatial sensitivity than did our NH listeners. Indeed, many of our HI listeners had ITD discrimination thresholds similar in magnitude to the spatial separation of adjacent streams in the small ITD condition (see (Dai et al., 2018)). To the extent that listeners focus attention to different features based on their relative perceptual reliability, it makes sense that compared to NH listeners, our HI listeners rely more on pitch differences across the streams to maintain focus on the target melody. This likely explains why HI listeners, as a group, showed no significant alpha lateralization even for large ITD condition when averaging over the duration of the roughly 3-s-long trials, while NH listeners did.

Even though the HI listeners in the current task did not appear to maintain focus using spatial attention, as noted above, most performed the task above chance levels and thus used spatial cues at least initially. The failure of our HI listeners to show sustained alpha lateralization suggests that once the HI listeners (or perhaps even the NH listeners attending to sources separated by small ITDs) “latched on” to the target, they did not maintain attention using spatial focus, relying instead upon the pitch differences between sources and the regular timing of the notes within the target melody.

## Implications

Our results provide further evidence for alpha lateralization as a correlate of auditory spatial attention. For NH listeners who have good spatial acuity, sustained alpha lateralization over the duration of the melodies is significant when ITDs are large. While alpha lateralization reflects the use of sustained spatial attention in NH listeners hearing large spatial separations, no sustained alpha lateralization was observed in HI listeners. This lack of lateralization suggests that HI listeners do not (or cannot) rely on spatial cues to sustain attention, consistent with their poor ITD discrimination thresholds. This inability to use spatial cues as effectively as do NH listeners helps explain the overall poorer performance of our HI listeners.

HI listeners often report difficulty communicating in noisy environments, contributing to a sense of social isolation in everyday settings. If it were possible to decode where or what an individual is trying to attend, then technology could be designed to assist object selection (e.g., by filtering out sound sources that a listener is trying to ignore). For instance, knowledge of where listeners are focusing spatial attention could be used to amplify sound from one direction while suppressing irrelevant sounds from others. EEG technology is being investigated for this purpose in many labs today (Choi et al., 2013; Eyndhoven et al., 2017; O’Sullivan et al., 2017), as it is relatively low cost, portable, and noninvasive. If EEG correlates of attentional focus could be reliably decoded, they could be used to create smart hearing devices that help a listener switch and maintain attention as needed.

Our previous analysis showed that HI listeners are less effective at modulating N1 ERPs than are NH listeners (Dai et al., 2018), calling into question the potential utility of using these neural correlates of attention in next-generation listening devices for the listeners most in need of such aid. Here, we find that alpha lateralization signatures for where a listener is attending are also weak in HI listeners, suggesting HI listeners do not rely on sustained spatial attention in conditions where NH listeners do. Importantly, however, it may be that HI listeners do not rely on sustaining spatial attention because other features are more reliable. If a next-generation hearing device correctly determined where a listener was trying to focus spatial attention and modified the sound entering their ears effectively, HI listeners might learn to rely on spatial attention. Thus, even though the HI listeners do not show strong, robust neural correlates of spatial attention, there is still the possibility that they could be trained to use such neural signals to control an effective device. Future work thus may require not only building devices that look for typical neural correlates of attentional control, but training listeners to engage listening strategies that their past experience has taught them to avoid.

## Acknowledgments

This work was supported by the Oticon Foundation and the National Institute for Deafness and Communication Disorders (Grants DC01598 and DC015760). This work was also supported by NIH Award No. 1RO1DC013825.

